# Tissue-specific modification of cellular bioelectrical activities using the chemogenetic tool, DREADD, in zebrafish

**DOI:** 10.1101/2021.06.22.449481

**Authors:** Martin R. Silic, GuangJun Zhang

## Abstract

Cellular electronic activity plays an essential role in neuronal communication. Manipulation and visualization of cellular membrane potential remain essential tasks in order to study electrical signaling in living organisms. Light-controlled optogenetic and designed chemical-controlled chemogenetic tools were developed to manipulate cellular electric activities for neuroscience research. One of the most common chemogenetic tools is DREADD (designer receptors exclusively activated by designer drugs). It has been extensively utilized due to its convenience and long-lasting effects in murine and primate models, but not in zebrafish, a leading model organism in various research fields. Here, we first establish multiple tissue-specific transgenic zebrafish lines that express two different DREADDs with a genetically encoded voltage indicator, ASAP2s. We observed voltage changes in zebrafish melanophores, epidermis, and neurons by hM4DGi or rM3DGs receptors measured by ASAP2s fluorescence intensity. Alteration to melanophore bioelectricity by DREADD generated dynamic electric signals and resulted in morphological alterations to pigment cells. We also tested a few agonists and found that the latest generation performs better than clozapine N-oxide (CNO). Collectively, our experiments demonstrate that DREADD can be utilized to manipulate cell-specific membrane potential in the zebrafish model. The availability of this tool in zebrafish will offer a new resource for a variety of bioelectricity research fields such as neuroscience, cardiology, and developmental biology.

## INTRODUCTION

Cellular bioelectric signaling has been extensively investigated in neuromuscular excitable cells due to the important roles of action potential signals and resting membrane potential (1). Recently, accumulating evidence reveals that cellular electric signaling is also an important player for regulating hormone release, embryonic development, wound healing, and regeneration (2,3). Non-invasive perturbation and visualization of cellular electrical activity in real-time and *in vivo* are the central requirements for studying cellular electric signaling (4).

To meet the rapid growth of neuroscience, genetically encoded tools have been developed for perturbation and visualization of cellular bioelectrical activity (5). Channelrhodopsin-based optogenetic tools can enhance or repress neuronal firing on a millisecond scale, and have been successfully applied to elucidate neuron type, activity, circuits, and behaviors (6,7). However, optogenetics generally requires complicated equipment, constrained animals free of movement, and can only modify neuronal activity in the short term (second-minutes). The chemogenetic tools were then developed to meet these needs. Chemogenetics use synthesized small molecules to activate engineered proteins (channels or receptors), modifying cellular electricity over a relatively long period (4,8). DREADDs (Designer receptors exclusively activated by designer drugs) are one group of the most commonly used chemogenetic tools (9,10). The DREADDs are composed of four tools (hM3DGq, rM3DGs, hM4DGi, and KORD) based on different mutated genetically engineered muscarinic receptors. Depending on the downstream G protein-coupled receptor signaling, the DREADD tools can modify cellular bioelectric activity bidirectionally. For example, hM3DGq enhances neuronal excitability, while M4DGi and KORD inhibit cellular excitability. DREADDs have been successfully used to elucidate behavior, circadian disorders, pathways related to cognitive impairments, eating disorders, neuronal plasticity, memory, and more in various animal models (11-14). Furthermore, improvements have been made to increase the selection of ligands with improved specificity and affinity (15-17).

Genetically encoded tools for visualizing and measuring cellular electrical activity *in vivo* are equally important for studying cellular electricity. Revolutionary biosensor tools, genetically encoded voltage indicators (GEVIs), for measuring cell membrane voltage have been developed for neuroscience (18-20). These GEVIs measure cellular voltage based on either FRET (fluorescence resonance energy transfer) or levels of fluorescence intensity. The advantages over traditional electrical physiology recording include non-invasive, real-time, and nanosecond sensitivity. Among them, the ASAP1-3 (Accelerated Sensor of Action Potentials) have been applied to a variety of model organisms such as fuit fly, mouse, and zebrafish (21-25).

Zebrafish have extensively been used for studying embryonic development and modeling human diseases, including cancer. This is because of the many advantages such as vertebrate biology, tractable genetics, external development, and early embryo transparency (26,27). We and others demonstrated that ASAP1 reported embryonic cellular voltage changes and neural activities in zebrafish (24,28), and a few optogenetic tools were just successfully adopted to the zebrafish model (29). However, DREADDs have been tested and reported to be non-functional in zebrafish (30). Thus, there is still a critical need for a chemogenetic tool that can modify zebrafish cellular electricity in the long term.

Here, we generated DREADD transgenic zebrafish lines and tested their function in zebrafish embryos and larvae using newly developed agonists. We demonstrated that this chemogenetic tool is functional in melanophore, neuron, and epithelial cells. Thus, the DREADD tools and our transgenic zebrafish lines can be an excellent resource for the zebrafish community for investigating cellular electricity.

## RESULTS

### Developing transgenic zebrafish lines to express DREADDs

Chemogenetic tools have been demonstrated successfully in neuronal studies with murine and primate models. However, the chemogenetic tools, both DREADD and PSAM (pharmacologically selective effector molecules), were found infective in injected zebrafish embryos (30). Recently, the PSAM tool was reported functional in zebrafish using the Tol2 transposon-based transgenic approach (31). Thus, we reasoned that the DREADD tool might also work in transgenic zebrafish. We then created melanophore-specific transgenic zebrafish lines to co-express hM4DGi and ASAP2s using the *mitfa2*.*1* promoter (Fig. 1A). This fish will allow us to simultaneously examine bioelectric changes during the process of cell membrane voltage manipulation. Adapting to investigate bioelectricity in multiple tissues, we also take advantage of the Gal4-UAS (Upstream activator sequence) artificial binary gene expression system (Fig.1B) (32). We made zebrafish lines in which a UAS promoter drives either hM4DGi or rM3DGs, together with ASAP2s. Additionally, we made the melanophore (*mitfa2*.*1* promoter) and basal epithelial cell (*tp63* promoter) specific Gal4FF fish lines (Fig.1C). We chose these two cell types because they are on the surface of the fish embryos, where the agonist chemicals can reach the cells easily, and it is convenient for us to image the cell membrane voltage change. These fish lines allow us to assess the effects of DREADD with different agonists (Fig. 1D). All the transgenic fish were outcrossed with wild type to F_1_ or F_2_ to clean the genetic background before use for experiments.

**Fig. 1.**
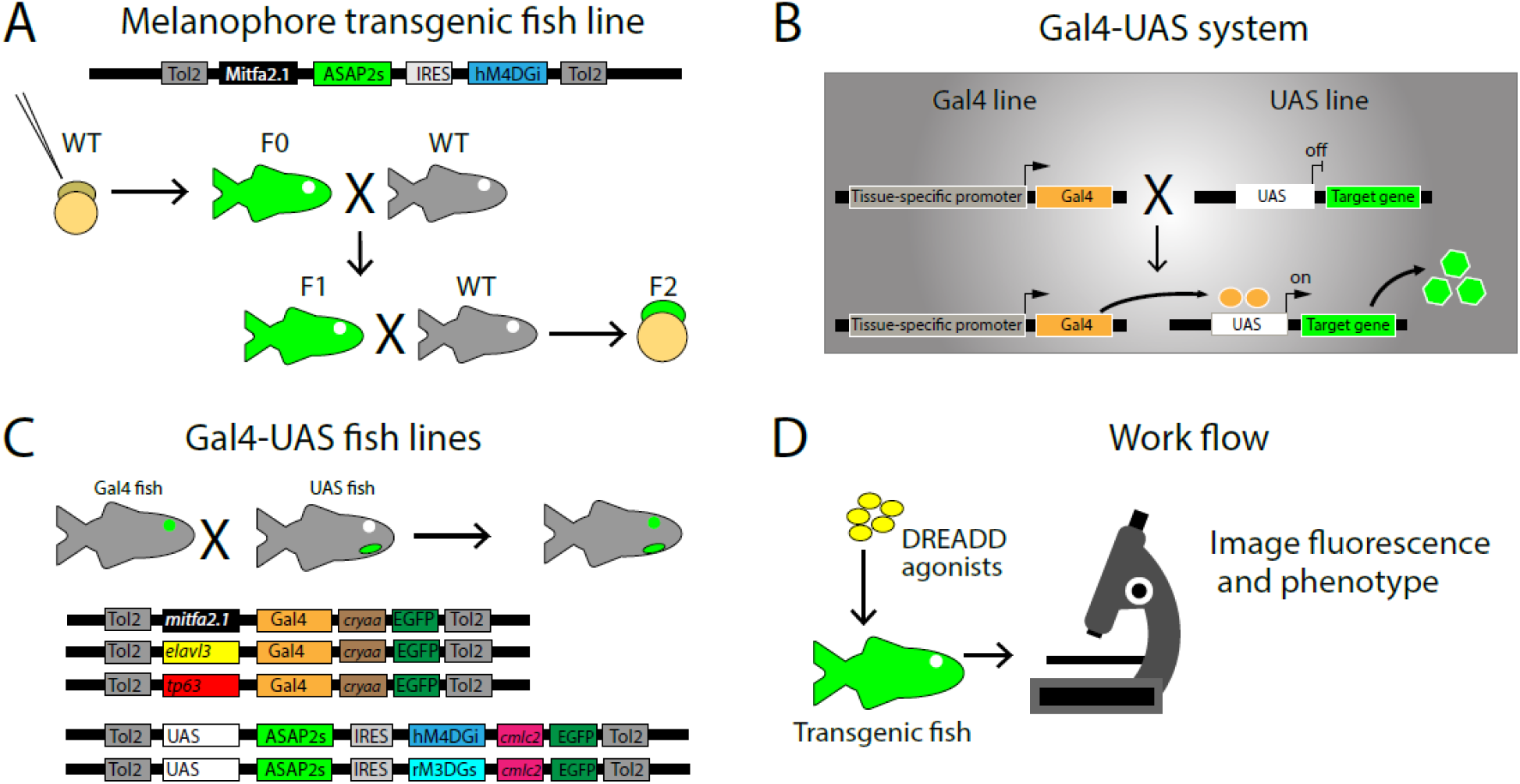
Illustration of DREADD transgenic fish lines and experimental workflow. **A**. Tol2 construct and method to produce stable DREADD zebrafish line, Tg (*mitfa2*.*1*: *ASAP2s-IRES-hM4DGi*). X, fish cross. Black arrow, fish raising or produced. **B**. Illustration of principles of the Gal4-UAS system. **C**. Diagrams of the Tol2 transposon plasmid constructs used for the Gal4-UAS transgenic zebrafish lines. The Gal4 fish lines have an eye maker (green), *cryaa:EGFP*. The UAS fish lines have a heart marker (green), *cmlc2:EGFP*. X, fish cross. Black arrow, fish raising or produced. Tol2, Tol2 transposon minimal flanking DNA sequences. IRES, internal ribosome entry site. **D**. Basic workflow: DREADD agonists were used to treating transgenic DREADD zebrafish embryos to cause cell bioelectric changes, then the fish embryos were subjected to green fluorescence imaging and quantification.

### DREADDs work in zebrafish embryos, evidenced by ASAP2s fluorescence intensity changes

Since ASAP2s, a sensitive cell membrane voltage reporter, was included in our transgenic zebrafish, we reasoned that the green fluorescence of fish embryos would change if the DREADD tool works. To test this, deschloroclozapine (DCZ), one of the most recently reported highly potent DREADD agonists, was tested with a relatively high dosage by adding 1µL of 100mM DCZ to an imaging slide with about 400µL of fish system water (∼250 µM). We treated 2dpf (day post-fertilization) Tg(*mitfa2*.*1:ASAP2s-IRES-hM4DGi*) fish embryos and indeed detected increased fluorescence in the melanophores on the top of the head about 5-10 minutes after treatment (Fig. 2A, D). This increased fluorescence is consistent with the hyperpolarizing activity of hM4DGi. Next, we examined epithelial cells using the fish embryos from Tg(*tp63:Gal4VP16;cryaa:EGFP*) and Tg(*4xnrUAS:ASAP2s-IRES-hM4DGi; cmlc2:EGFP*) fish cross. Similarly, the DCZ treatment resulted in fluorescence intensity increases of the epithelial cells in the head region (Fig. 2B, E) and caudal fin folds (Fig. 2bb, ee). To further test hM4DGi’s function in neurons, we first injected the *elavl3* promoter-driven Gal4FF construct into Tg(*4xnrUAS:ASAP2s-IRES-hM4DGi;cmlc2:EGFP*) fish embryos and raised them to 2dpf. Then, we treated the fish embryos with DCZ. As expected, we found the neurons in the neural tube showed enhanced green fluorescence in treated fish embryos (Fig. 2C, F). Thus, the inhibitory DREADD, hM4DGi, is indeed able to induce hyperpolarization within zebrafish. To further test the DREADD tools in zebrafish, we examined the excitable DREADD, rM3DGs. Both melanophores (Fig. 2G, J) and neurons (Fig. 2I, L) showed decreased green fluorescence after DCZ treatment. This decreased green fluorescence is consistent with depolarizing activity of rM3DGs. In contrast, the epithelial cells showed increased green fluorescence (Fig. 2H, K). This unexpected opposite result may be caused by the epithelial cell’s physiological response for maintaining its resting membrane potential. Overall, our experiments demonstrated that DCZ could activate both hM4DGi and rM3DGs in zebrafish.

**Fig. 2.**
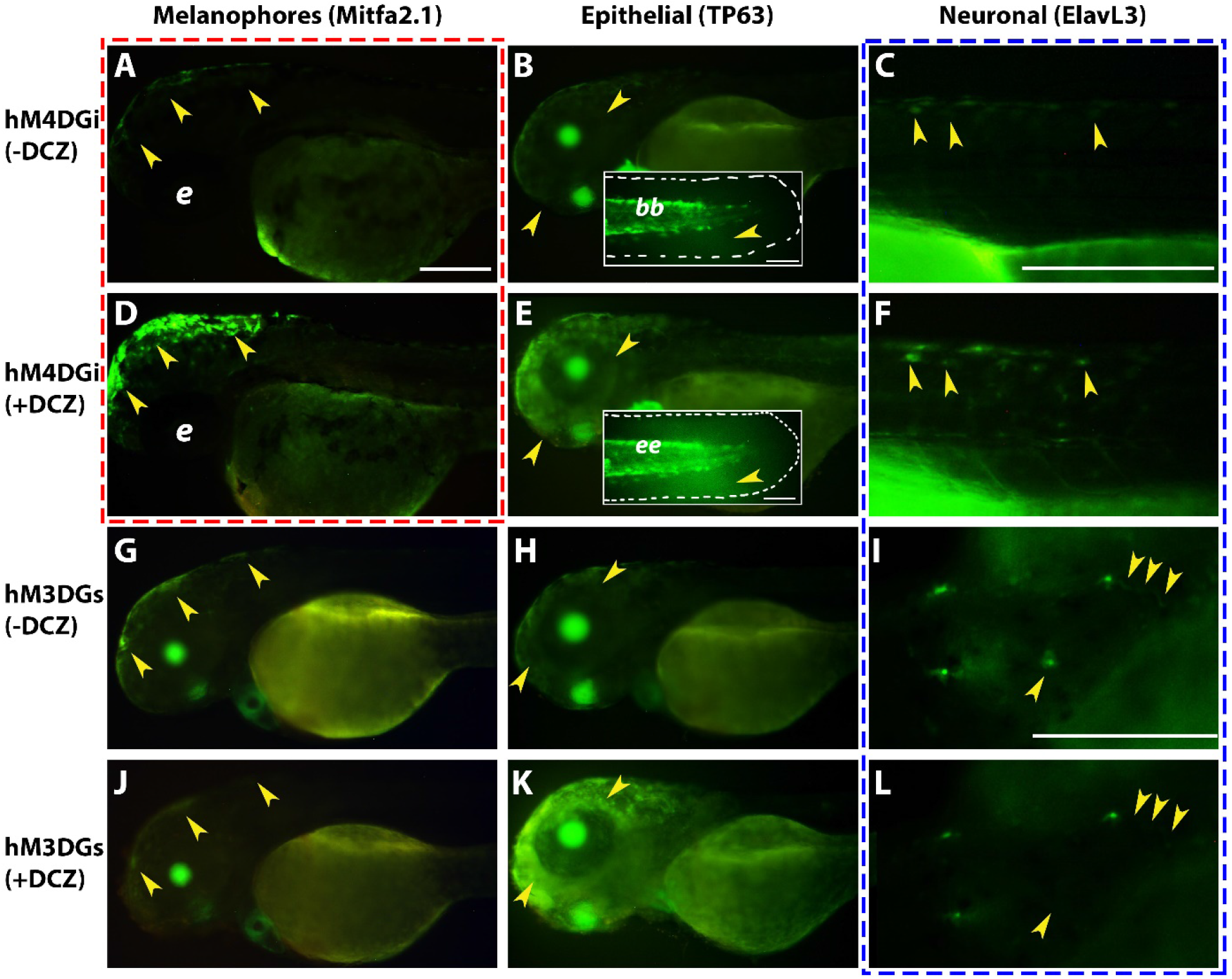
Cell membrane voltage manipulation by DREADD in zebrafish embryos. **A-F**. Transgenic zebrafish expressing ASAP2s-IRES-hM4DGi in melanophore (*mitfa2*.*1*), basal epithelial cell (*tp63*), or neuron (*elavl3*) promoter, respectively. **A-C**. Fish embryos before DCZ treatment. **D-F**. The same fish corresponding to the **A-C** panels after treatment with DCZ. Inserts **bb** & **ee:** caudal fin images of Tg(*tp63:ASAP2s-IRES-hM4DGi*) fish before and after DCZ treatment, respectively. Yellow arrows indicate specific cells with increased levels of fluorescence. **G-L**. Transgenic zebrafish expressing ASAP2s-IRES-hM3DGs in melanophore (*mitfa2*.*1*), basal epithelial cell (*tp63*), or neuron (*elavl3*) promoter, respectively. **G-I**. Fish embryos before DCZ treatment. **J-L**. The same fish corresponding to the **G-I** panels after treatment of DCZ. Panels (**A, D**) with red dotted lines are from a cross between the Tg(*mitfa2*.*1*:ASAP2s-*IRES-hM4DGi*) and wild type. Panels (**C, F, I, L**) with blue dotted lines were from Tg(*UAS:ASAP2s-IRES-hM4DGi*) or Tg(*UAS:ASAP2s-IRES-hM3DGs*) fish injected with *elavl3*:Gal4FF plasmid construct. The remaining panels (**B, E, G-H, J-K**) were offspring from crosses of Gal4 and UAS transgenic fish lines. All the fish embryos are two days old. Yellow arrows indicate specific cells with altered levels of fluorescence. Only matching (before and after DCZ treated) embryos are directly comparable for fluorescence intensity levels (**A** & **D, B** & **E, C** & **F, G** & **J, H** & **K, I** & **L**.) Scale bars = 250 μm except for panel inserts **bb** and **ee** where scale bar = 50 μm.

### DREADD activation causes dynamic bioelectric changes in zebrafish melanophores

We have demonstrated that DCZ can activate *hM4DGi* in the Tg(*mitfa2*.*1:ASAP2s-IRES-hM4DGi*) fish causing fluorescence to increase after 5-10 minutes. To further examine the exact bioelectric changes that take place after hM4DGi activation leading up to overall fluorescence increase, we decided to record treated larvae with time-lapse imaging immediately. We found that the green fluorescence intensity increase was not linear. Instead, the fluorescence intensity fluctuated, but the overall intensity increased with time extended. In addition, the fluorescence intensity of adjacent melanophores also fluctuated (Fig. 3A-G). These results indicate a melanophore membrane potential homeostasis, which may take time to change using DREADD and its agonist. These results also might indicate why all DREADD expressing cells do not immediately show uniform fluorescence change.

**Fig. 3.**
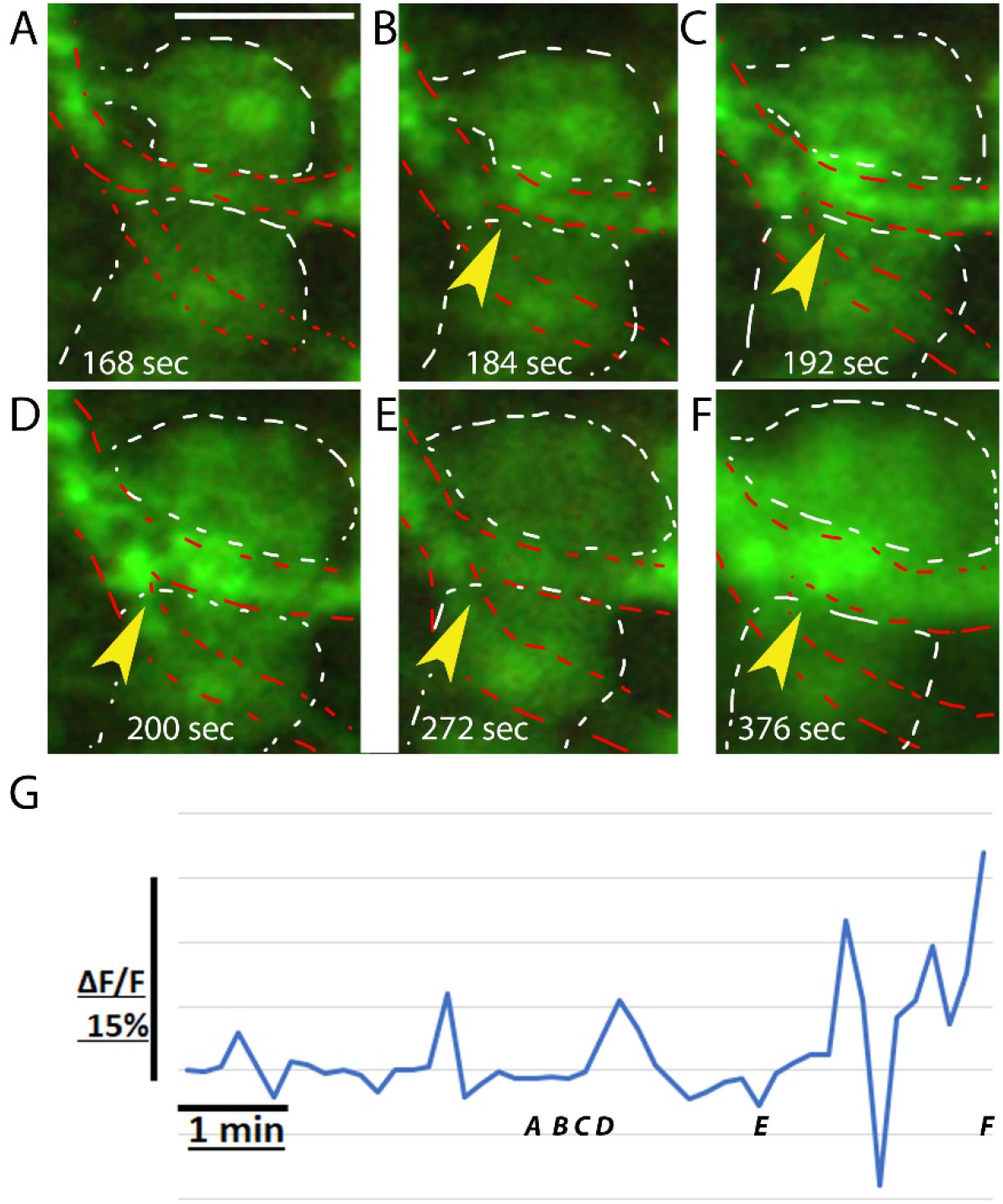
Melanophores show dynamic fluorescence changes after DREADD modification. **A-F**. Six different time points of melanophore imaging in the head region of a 2dpf Tg(*mitfa2*.*1*:ASAP2s-*IRES-hM4DGi*) zebrafish embryo after DREADD activation. The time scale is in seconds. Arrows point to changes in GFP intensity in the same location. **G**. ΔF/F quantification of melanophore fluorescence intensity changes over a 6-minute duration. ASAP2s fluorescence shows fluctuations in intensity. Scale bar = 25 μm. The corresponding time-lapse video can be found in the Supplementary file S1.

### New generation DREADD agonists are more potent and specific

The original DREADD agonist, Clozapine-N-oxide (CNO), was found not as ideal due to its back-conversion to clozapine, an activating molecule with psychoactive properties prescribed to treat schizophrenia. This can have adverse side effects on studies linked to animal behaviors in mouse studies (33). More potent agonists were screened out to overcome this issue. To select the best agonists for DREADDs in zebrafish, we tested Compound 21 (C21) (34), JHU37160 (15), and CNO, then compared them to DCZ. One day old fish embryos were collected from Tg(*mitfa2*.*1:ASAP2s-IRES-hM4DGi*) fish in-cross and were averaged into four wells in a 6 or 12-well plate. Next, the fish embryos were treated with the four agonists at various concentrations according to previously reported working concentrations for the mammalian system. Twenty-four hours later, the brightest embryos from each well were sorted out and imaged under the same microscope settings and compared to untreated embryos (Fig. 4A-B). Fish embryo images were processed for calculating the corrected total cell fluorescence (CTCF), which is a convenient estimate of fluorescence intensity that takes into account area and background fluorescence. Then, ASAP2s green fluorescence intensities were compared at each concentration of the agonists. For all the agonists, higher concentration resulted in more significant fluorescence changes (Fig. 4C). However, the effective dosage that leads to fluorescence increase varied greatly. CNO and C21 performed the weakest, while the latest generation, JHU37160, and DCZ, had the best response at a dramatically lower dosage. The potency of these agonists in zebrafish is consistent with recent reports in murine systems.

**Fig. 4.**
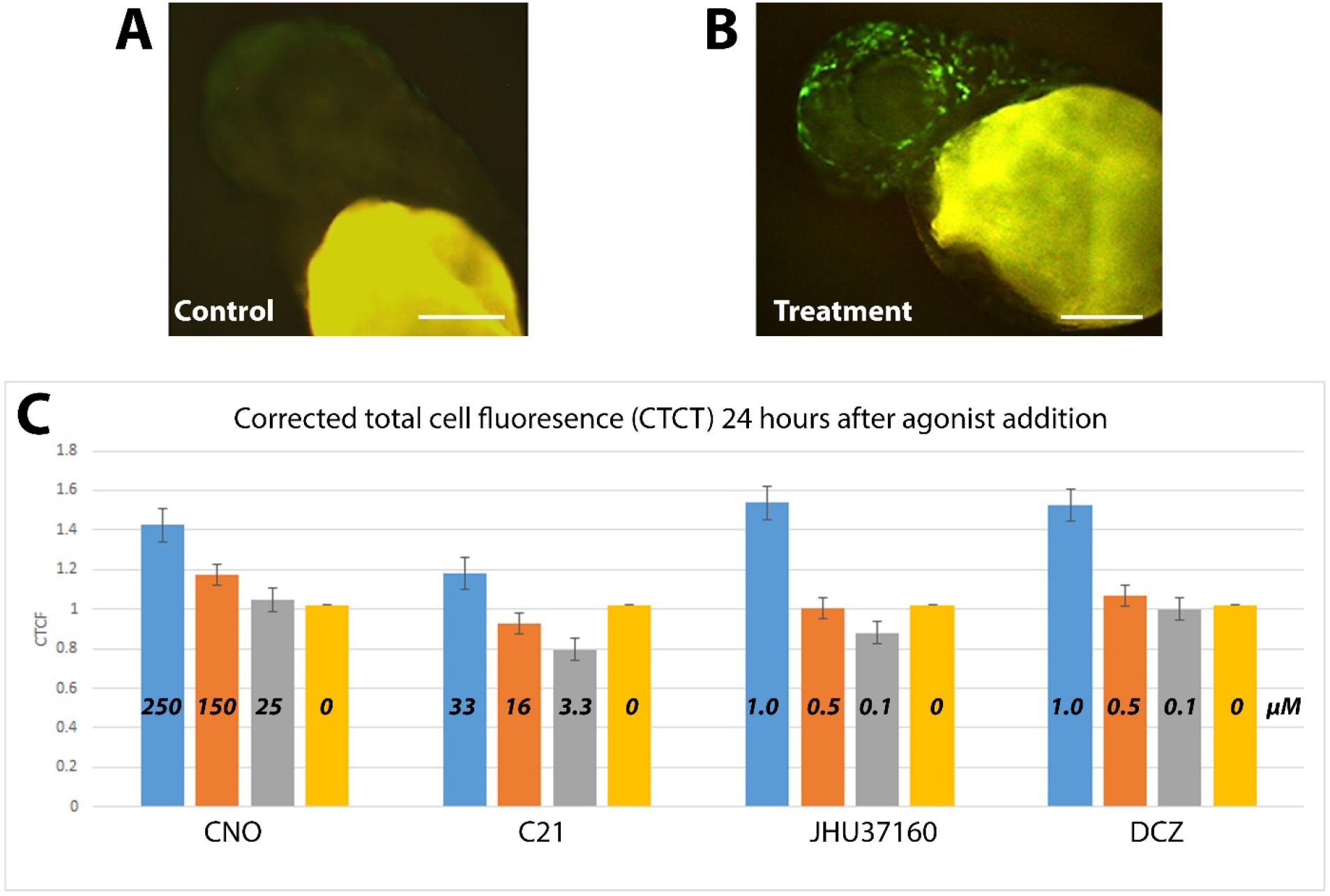
New DREADD agonists, DCZ, and JHU37160 have higher efficacy. **A**. A representative embryo from the Tg(*mitfa2*.*1*:ASAP2s-*IRES-hM4DGi*) zebrafish. The melanophores in the head region display minimal green fluorescence. **B**. A representative embryo shows bright pigment fluorescence after agonist treatment. **C**. Quantification of corrected total cell fluorescence (CTCF) after 24-hour treatment with different concentrations of CNO, C21, JHU37160, and DCZ. The concentrations (μM) are labeled in the vertical bars with different colors. All the fish embryos are two days old. Scale bars = 250 μm.

### DREADD functional validation by melanophore morphological changes

We successfully modified cell membrane potential that can be measured via the ASAP2s voltage reporter. Whether this DREADD induced voltage change is enough to effect the *in vivo* biology remains unknown. To address this question, we treated Tg(*mitfa2*.*1:ASAP2s-IRES-hM4DGi*) fish larva with 20uM DCZ at 2dpf then raised them for five days (2dpf-7dpf). We found that the treated fish larvae developed hyperpigmentation compared to the untreated sibling control group (Fig. 5A-B). To figure out whether this melanophore hyperpigmentation was caused by an increased number of melanophores or melanophore dispersion, we treated these fish larva with 1mM epinephrine (α_2_-adrenoceptor agonist), which is known to cause melanosome aggregation. Epinephrine caused pigment granule contraction in DCZ treated fish larvae (Fig. 5C-D). Interestingly, embryos treated with epinephrine that were re-treated with DCZ caused pigments to start expanding (Fig 5E). To rule out the possible side effect of DCZ and examine the efficacy of other agonists, we did the same experiment with the other three agonists. The DCZ and JHU37160 agonists have a higher percentage of hyperpigmentated larvae, and a majority of them (81% DCZ and 72.7% JHU37160) are transgenic fish confirmed by post-experimental genotyping of a portion of treated embryos (Table 1). In contrast, the CNO and C21 agonist treatment yielded fewer pigment responses. All the agonist treatments produced a hyperpigmentation phenotype, but DCZ had the lowest off-target rate, suggesting this is a good choice for zebrafish experiments. To test the response of metamorphic melanophores to DREADD induced bioelectric changes, Tg larvae were raised to 2 weeks of development (∼6.5mm), and then treated with DCZ (20 μM) for 1 week every 2 days. Fish were then imaged at about 30 days of development (9-10mm) for changes to pigmentation. Comparable-sized untreated transgenic fish were raised as a control. We found that one fish developed an expanded melanophore pigmentation on the lateral sides and fins compared to untreated controls (Fig. 5F-G). In summary, our results confirmed that the DREADD tools indeed are functional and can be used for relevant biological studies.

**Table 1.** Zebrafish larva pigmentation change is caused by different DREADD agonists.

| Larval genetic background | DCZ | JHU37160 | CNO | C21 |
| --- | --- | --- | --- | --- |
| Mixed transgenic and wildtype siblings with pigmentation change | 49.20% | 60.10% | 39.00% | 8.60% |
| Genotype confirmed positive transgenic embryos with pigment change | 81.00% | 72.70% | 60% | 50% |
| Wildtype embryos treated with pigmentation change | 5.50% | 17.90% | 12.50% | 5% |
| Untreated transgenic embryos with pigmentation change | 1.25% | 3.75% | 2.50% | 1.25% |

**Fig. 5.**
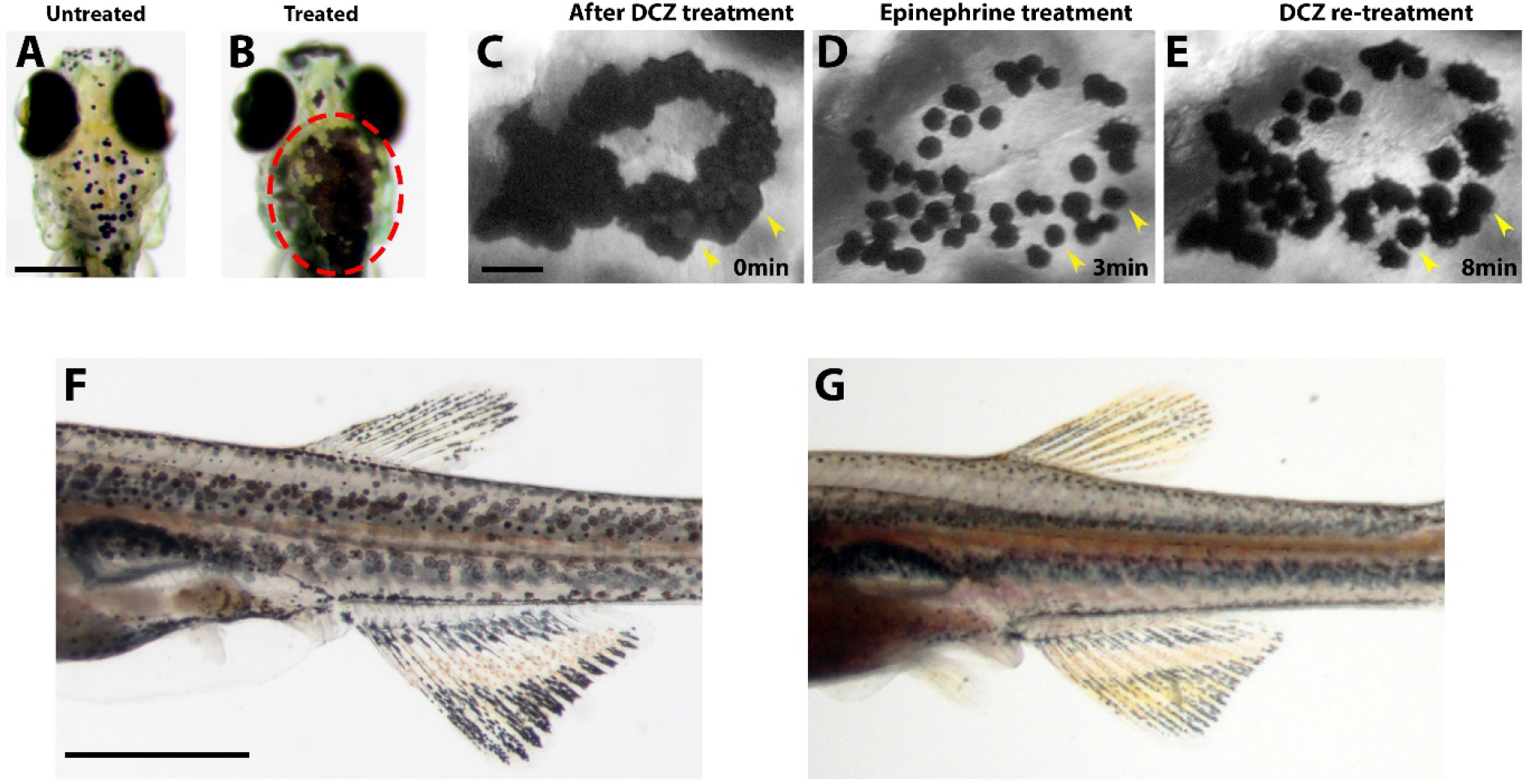
The hM4DGi receptor activation causes hyperpigmentation in 1-week old and metamorphic fish larvae. **A**. A representative untreated 1-week old fish larva. **B**. A representative treated fish larva with hyperpigmentation (circled with red dotted line). **C**. Dorsal view of another Tg(*mitfa2*.*1*:ASAP2s-*IRES-hM4DGi*) fish larva that was treated with DCZ. The melanophores are dispersed. **D**. The same fish larva imaged 3 minutes after treatment with 1mM epinephrine. The melanophores are aggregated. **E**. The same epinephrine-treated larva was re-treated with DCZ. The image was taken 8 minutes after treatment. The melanophores dispersed again. Yellow arrows point to melanophores that are dispersed, contracted, and re-dispersed, respectively. **F**. 9.4mm fish that was treated with DCZ for one week (20 μM) every two days starting at about 6.5mm of development. Melanophores appear to be dispersed over a larger area. **G**. Comparable sized untreated transgenic fish. Melanophores appear smaller and more uniform. Scale bar = 250 μm for **A-B** and 50 μm for **C-E**.

## DISCUSSION

The chemogenetic tools are useful for manipulating bioelectricity and have been demonstrated successfully in neuroscience with murine and primate models. DREADD and PSAM tool adaptation was attempted but not successful in zebrafish (30). Recently, we showed that the bioelectricity of the somites is involved in fin patterning (35). To further investigate the roles of bioelectricity during embryonic development, there is a need for chemogenetic tools which allow us to manipulate bioelectricity for days and months. In addition, we have successfully adapted the ASAP1 voltage sensor to zebrafish using the Tol2 transposon-based transgenic approach (24). Recently, the PSAM tools were demonstrated functional in zebrafish using the same transgenic approach (31). This motivated us to re-examine the possibility of adopting DREADDs in zebrafish.

There are a few caveat issues that could make a difference between our work and the previous study. First, the sensitivity of the detection method may be a key factor. We use the newly developed ASAP2s, which was reported to have a high level of sensitivity to cellular membrane voltage changes. In comparison, locomotor activity was used for measuring the bioelectric changes previously. It is possible that there is some level of bioelectric change, but not enough to drive the locomotor activity change. Second, we examined the DREADDs using transgenic zebrafish lines, not the direct injection as in the former study. The transgenic fish provide stable and specific DREADD expression in the fish embryos for the targeted tissue. Thus, it reduces many stochastic expressions for measuring electric activities. In contrast, the direct injection of DREADD constructs will yield different levels of expression in many places. Third, we tested a few agonists in our current study and found the formerly used CNO less potent than the newly developed agonists. In our experiments, CNO was able to cause DREADD activation and induce morphological melanophore changes. Still, these results were not as strong and less specific compared to the latest generation DREADD ligands DCZ and JHU37160. The use of CNO could impact the assessment dramatically, especially in the situation of less tissue-specifically expressed DREADDs. When we injected the *elavl3*:Gal4FF plasmid construct into Tg (*UAS*:DREADD-ASAP2s) transgenic fish embryos, only a few larval neurons were observed with fluorescence changes. One solution is to improve the DREADD expression levels. In the future, zebrafish fish-codon optimization, utilizing the zebrafish Kozak sequence before an ATG start codon, and adding dORF (downstream open reading frame) could be attempted for this purpose (36,37). Lastly, the examination time after treatment might also affect the judgment. In our experimental system, a higher concentration of ligands is still needed in order to visualize more impressive fluorescence changes over a shorter period of time (∼5minutes). This could be caused by tissue penetration, fish metabolism, or chemical potency.

There are numerous zebrafish pigment pattern mutants that affect proteins that regulate charged molecules, such as ion channels and gap junctions (38). Alteration of these ionic regulators can cause morphological changes to zebrafish pigments and disrupt normal stripe formation. It was previously reported that melanophore cell membrane voltage manipulation with the optogenetic tool ChR2 was able to disrupt stripe formation in metamorphic and adult zebrafish (39). Here, we chose zebrafish melanophores as a model and tested two DREADDs, hM4DGi and rM3DGs, and four agonists. We demonstrated that both DREADDs are functional to change cell membrane voltage measured by ASAP2s. These voltage alterations were validated in basal epithelial cells and neurons. Moreover, we were able to generate a melanophore hyperpigmentation phenotype via hM4DGi and DCZ in 1-week larvae, confirming the biological functions of DREADD in zebrafish. hM4DGi activation of metamorphic melanophores also results in hyperpigmentation compared to untreated controls. These results provide additional validation for the biological impact, as well as the potential use of zebrafish DREADDs in older fish over longer periods of time.

In summary, we generated tissue-specific DREADD transgenic zebrafish lines and tested their function in zebrafish embryos and larvae using different agonists. We demonstrated that this chemogenetic tool works in zebrafish melanophores, neurons, and epithelial cells. The newly developed agonists are much more potent and specific compared to CNO. We expect the DREADD tools and our transgenic fish lines will meet the critical need for neuronal and embryonic bioelectric studies. Moreover, they can be a great resource to the zebrafish community.

## MATERIALS AND METHODS

Zebrafish were raised and maintained at the Purdue animal housing facility following Association for Assessment and Accreditation of Laboratory Animal Care (AAALAC) approved standards. Experiments were carried out according to Purdue Animal Care and Use Committee (PACUC) approved protocols. All zebrafish experiments were carried out in wild-type TAB fish. Zebrafish were maintained according to the zebrafish book. Zebrafish embryos were staged based on the Kimmel staging guide.

### Tol2 constructs, microinjection, and zebrafish transgenic lines

Tol2 transposon plasmids were generated using three-fragment Gateway cloning-based Tol2 kit (40).The 5’ end entry plasmids p5E-*mitfa2*.*1 (plasmid #81234)*, p5E-*elavl3* (plasmid #72640) (41), p5E-4xrnUAS (plasmid #61372) (42) were acquired through Addgene. pME-Gal4FF were subcloned from pCR8GW-Gal4-VP16-FRT-kan-FRT (43), a gift from Dr. Koichi Kawakami. p5E-tp63 was a gift from Dr. Qing Deng (44). The pENTR-D-ASAP2s plasmids were generated by site-directed mutagenesis from pENTR-D-ASAP1 (24) using the primers (ASAP2s-F: ATA TTT CAG CTG GCT TCA CAG AAG AAA CAA CTT GAA GTG G and ASAP2s-R: AGC CAG CTG AAA TAT TCT TAT TAA GAT AAC AAT TCT CAG AAC TCG AAG AAG AG). The p3E-*IRES-hM4DGi* and p3E-*IRES-rM3DGs* were generated by subcloning from pAAV-hSyn-DIO-HA-hM4DGi-IRES-mCitrine (plasmid# 50455) and pAAV-hSyn-DIO-HA-rM3DGs-IRES-mCitrine (pasmid# 50456) into p3E-IRES-EGFP vector (Tol2 kit #389), respectively. All the subcloning and site-directed mutagenesis were performed using the In-Fusion cloning system (Takara Bio). Final entry constructs were verified by Sanger sequencing. The final constructs were built using LR Clonase II Plus enzyme (Thermo Scientific) following the manufacturer’s instructions.

pCS-zT2Tp plasmid (a gift from Koichi Kawakami) was used as a template to generate Tol2 transposase messenger RNA (mRNA) using the mMESSAGE mMACHINE SP6 transcription kit (Thermo Scientific). Capped and tailed mRNAs were purified by Monarch RNA Cleanup Kit (NEB) according to the manufactory guide and eluted in DEPC-treated water. Microinjection of Tol2 expression constructs (25 ng/µl) and Tol2 mRNA (50 ng/µl) with 0.025% phenol red (P0290; Sigma) was performed on one-cell-stage zebrafish embryos under a dissection microscope (SMZ445; Nikon, Garden City, NY) using a PV820 pneumatic PicoPump (World Precision Instruments). About ten injected embryos were sampled for gDNA isolation using the HotSHOT method after at least 8 hours (45). Tol2 transposon efficiency was verified with an excise assay (46). Once the excise assays confirmed the Tol2 activity, the remainder of the injected fish embryos were raised to adulthood. TAB wild-type fish were crossed with F_0_-injected adult fish, and positive fluorescent transgenic zebrafish embryos were selected and raised to adults as stable F_1_ transgenic fish lines.

### DREADD ligand addition and fluorescence imaging

To detect cell membrane potential changes via ASAP2s imaging, embryos were raised to 2 days post fertilization (dpf), then anesthetized in a 0.05% Tricane solution. For fluorescence imaging experiments, embryos were raised in fish system water without methylene blue to reduce autofluorescence and treated with 1X PTU (1-phenyl 2-thiourea, 0.2mM) to reduce pigments. Anesthetized embryos were placed on a glass slide with fish system water and positioned properly for imaging. In order to visualize large changes in fluorescence intensity, 1µL of 100mM DCZ was added to slides containing Tg fish and ∼400µL of fish water for a final concentration of 250 µM. Untreated images were taken before agonist addition. After treatment, embryos were either monitored for 5-15 minutes for fluorescence changes and then imaged, or immediately recorded with time-lapse imaging. Exposure was set at the start of imaging (between 4000 and 8000 ms) and kept consistent for the entire length of time the same individual larva was imaged in order to compare levels of fluorescence intensity between images directly. All the DREADD agonists were purchased from HelloBio and diluted in water as 100mM stock solutions that were stored in a -20C freezer before use. Described concentrations of DREADD agonist were then added to the fish system water on the slide. Embryos were monitored for positional shifts and adjusted if they drifted from their original location. Next, time-lapse imaging was continued for 5-15 minutes. Cellular ΔF/F=((F_t_ – F_0_)/F_0_) was quantified using ImageJ to define a region of interest for mean fluorescence intensity at each time point. F_t_ is the fluorescent value at a given time t, and F_0_ is the starting fluorescence value. All images were acquired using Zeiss Axio CamMRc camera on Stereo Discovery.V12 or Axio Imager 2 compound microscope. For *elavl3*:Gal4FF, Tol2 plasmid was injected into 1-cell stage Tg (*UAS*:DREADD-ASAP2s) transgenic fish embryos. UAS fish lines were prepared for one-cell-stage microinjection as described above. Injected embryos were monitored and raised to 2dpf. Embryos with positive fluorescence, via GFP eye and heart markers, were sorted and used for agonist addition and fluorescence imaging, as previously mentioned. Unless specified otherwise, this type of signal is expressed as a fractional change in fluorescence intensity ΔF/F0.

### Quantifying fluorescence intensity from different ligand concentrations

About 25 Tg (*mitfa2*.*1*: *ASAP2s-IRES-hM4DGi*) embryos were treated with designated concentrations of ligands at 1dpf in 12-well plates. The following day, two embryos with the brightest fluorescence in each well were imaged with the same exposure and magnification. Images were later processed for corrected total cell fluorescence (CTCF) by subtracting the mean background intensity multiplied by the area for the ROI from the integrated density of fish head fluorescence (integrated density - (ROI area* mean background intensity). Values were then normalized to 1 for the untreated samples.

### Tracking 1-week larvae for DREADD induced phenotype

To track pigmentation changes caused by DREADD treatments, we crossed Tg (*mitfa2*.*1*: *ASAP2s-IRES-hM4DGi*) with TAB fish. Dechorionated 2dpf embryos were separated into groups of about 40 and placed into 6-well plates containing fish water with 0.05% methylene blue. This mixed group of siblings contained wildtype and transgenic larvae. Next, the water was replaced with fresh water containing a designated concentration of DREADD agonists. Embryos were treated once at 2dpf and raised until 7dpf. At 7dpf, larvae in each well were counted to obtain the number with increased head pigmentation. This was repeated, and both trials later added to the total number of fish tracked for each DREADD agonist (n=128).

For genotyping a representative portion of DREADD treated embryos, 96-well plates were used to separate single embryos from Tg (*mitfa2*.*1*: *ASAP2s-IRES-hM4DGi*) crossed with wildtype fish. Dechorionated 2dpf embryos were separated and placed into individual wells containing fish water with methylene blue. This mixed group of siblings contained wildtype and transgenic larvae. Next, water was replaced with 200uL of fresh fish water containing designated amounts of DREADD ligand. These two-day-old fish embryos were treated with either DCZ, JHU37160, C21, or CNO (n=48 for each agonist), respectively. Each well was tracked until seven days to observe pigmentation changes. On the 7^th^ day, the number of embryos with increased head pigmentation was counted. All embryos were then harvested for genotype using 100mM NaOH (hotshot method). PCR primers (ASAP1-genoF: ATA TGA CCT ACT CCT TCT CTG ACC and ASAP1-genoR: AGG TTA AGG TGG TCA CCA GG) were used to amplify the ASAP2s transgene to validate transgenic mutant correlation with phenotypic changes. These percentages were caculated as a representative of a total of 128 tracked embryos for each agonist.

For examining the variation of melanophore morphology, 7dpf Larvae were treated with epinephrine (1mM) to determine any changes to total pigment count and assess pigment expansion. Larvae were placed on a glass slide with fish system water and imaged under regular epifluorescent light. For evaluating the impact of bioelectric changes on metamorphic fish melanophores, transgenic larvae that were about two weeks of development (∼6.5mm) were treated with DCZ (20 μM) for one week every two days. Once fish reached about 30 days old (9-10mm), they were imaged under a microscope for pigmentation changes. Comparable-sized untreated transgenic fish were also imaged as a control.

## ACKNOWLEDGMENTS

The research was supported by the National Institute of General Medical Sciences of the National Institutes of Health (R35 GM-124913) to G.Z. The content is solely the responsibility of the authors and does not necessarily represent the official views of the funding agents.

## AUTHOR CONTRIBUTIONS

G.Z. conceptualization; M.R.S. and G.Z. formal analysis; G.Z. funding acquisition; M.R.S. and G.Z. Investigation; G.Z. Supervision; M.R.S. and G.Z. writing, review, and editing.

